# “Prevalence of of ABO and Rhesus (Rh) Blood Groups in Eastern UP Population”

**DOI:** 10.1101/2020.02.26.965988

**Authors:** Pradeep Kumar, Vandana Rai

**Author notes:** **Corresponding Author:** Dr. Pradeep Kumar, Human Molecular Genetics Laboratory, Department of Biotechnolgy, VBS Purvanchal University, Jaunpur (U P)222001, India.

## Abstract

Approximately 300 different types of blood groups are identified so far, the ABO and Rh antigens are still the clinically most significant and genetically most polymorphic of all human blood group systems to date. A total of 200 unrelated individuals from Uttar Pradesh were studied for the phenotype and allele frequency distribution of ABO and Rh (D) blood groups. In total 200 samples analyzed, phenotype B blood type has the highest frequency 36.5% (n=73), followed by O (34.5%; n=69), A (20.5%; n=41) and AB (8.5%; n=17). The O, A and B frequencies were 0.5849, 0.1571 and 0.2580 respectively. The overall phenotypic frequencies of ABO blood groups were B>O>A>AB. The variation in phenotypic frequencies between male and female might be due to small sample size of male sample. The allelic frequency of Rh-negative was 0.2.

## Introduction

Blood groups are antigenic determinants on the surface of red cells, platelets and granulocytes, but the use of the term is often restricted to antigens on red blood cells. Blood groups are defined by antibodies, usually alloantibodies produced by individuals who lack the corresponding antigen. Some blood group antibodies, such as anti-A and anti-B, are present in the plasma of a person, whose red cells lack the corresponding antigen, but most blood group antibodies are formed only in response to antigen positive red cells as the result of transfusion or pregnancy such as Rh (Daniels, 2001). Nearly 300 blood grouping antigens have been reported, the ABO and Rh are recognized as the major and clinically significant blood group antigens (Bauer, 1982). Rhesus blood group system was the 4^th^ system discovered and yet it is second most important blood group from the point of view of transfusion (Molison ,1979).

The ABO locus is located on chromosome 9 at 9p34.1-q34.2 and encodes glycosyltransferases. And ABO gene has three alleles namely *IA*, *IB* and *i* determine blood groups. *IA* produces A antigen, *IB* produces B antigen whereas *i* produces neither. IA and IB are mutant alleles and show codominance with each other but both are dominant over the wild type allele *i* . The concept of wild type allele is based on the fact that the allele more frequent in a population is a wild type allele (Gardner et al, 2001). Rh antigens are determined by three pairs of closely linked allelic genes located on chromosome 1.

The ABO and Rh blood groups have been studied extensively for their involvement in incompatibility selection and their distribution are studied in almost all the races and populations of the world like-Nigeria (Enosolease and Bazuaye, 2008), Kenya (Lyko et al., 1992), Palestine (Alishtayeh et al., 1988), Iraq (Mohamad and Jaff, 2010), Sudan (Hassan,2010); Pakistan (Hameed et al., 2002; Anees and Mirza,2005), Bangladesh (Talukdar and Das,2010), Saudi Arab (Adullah,2010), Jordan (Hanania et al,2007), Iran (Boskabady et al, 2005), Nepal (Pramanik and Pramanik,2000), and India (Chakraborty, 2010; Kumar et al.,2010; Rai and Kumar, 2010; Deepa et al, 2011;Kumar and Rai,2012). The genotype frequencies for a particular gene(s) in a population depend on the gene frequency and the proportions of different alleles of a gene in a Mendelian (panmictic) population are known as gene frequency. Estimates of gene’s frequency provide very valuable information on the genetic similarity of different populations and to some extent on their ancestral genetic linkage, despite the cultural and religious differences of the two populations. Keeping this in view, the present study was designed to see frequency of ABO and Rh blood group antigens in UP population.

## Materials and method

The study was done in Uttar Pradesh (UP). Over three months period (December 2010 to February 2011), a total of 200 unrelated individuals of both genders from Jaunpur and Varanasi, Azamgarh and Mau districts of UP were selected. Blood samples were collected from each subject and relevant data were also collected after taking written consent. Confidentially, of the data were maintained. Each subject, who accepted to participate in the study, received two sheets, (consent form, and questionnaire). The first sheet was a declaration form for each participant that she/he understood the project/study well, the second sheet was, a questionnaire that included profile/demographic data of the participants. The blood samples were collected by finger prick with sterile lancet and cleaning the puncture site with 70% ethyl alcohol. A drop of monoclonal anti-A, anti-B, monoclonal/polyclonal anti-D (Span) was added to a drop of finger prick blood on clean slide and mixed well. Results of agglutination were recorded immediately for ABO blood groups and after 2 minutes in Rh(D) (Bhasin and Chahal, 1996). The gene frequencies for this system were calculated according to the method of Mourant et al. (1976).

## Results and Discussion

In total 200 samples analyzed, phenotype B blood type has the highest frequency 36.5% (n=73, followed by O (34.5%; n=69), A (20.5%; n=41) and AB (8.5%; n=17) (Table 1).The O, A and B frequencies were 0.5849, 0.1571 and 0.2580 respectively (Table 1. The overall phenotypic frequencies of ABO blood groups were B>O>A>AB. Total numbers of samples were categorized by gender-wise, 68 samples were of females and 132 of males (Table 2).In female samples 20 individuals have O blood group, 17 individuals have A blood group, 25 individuals have B blood group and 6 individuals have AB group. In 132 male samples, O, A, B and AB blood groups were found in 49, 24, 48 and 11 individuals respectively. In female samples the phenotypic frequencies were B> O> A>AB, whereas in male samples overall phenotypic frequencies were O> B> A>AB (Table 2). The variation in phenotypic frequencies between male and female might be due to small sample size of male sample. Out of total 200 samples, 192 (96%) samples were Rh-positive and 8 (4 %) were Rh-negative. The allelic frequency of Rh-negative was 0.2 (Table 3).

**Table 1.**
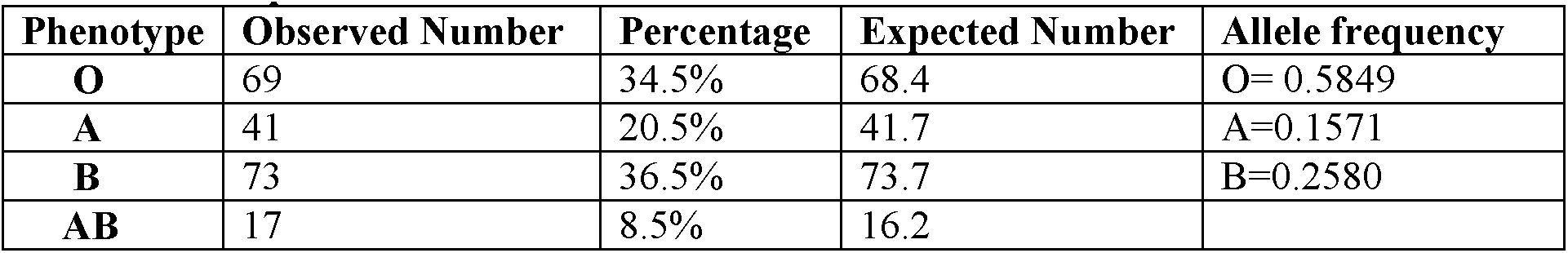
Distribution of the ABO blood group and their allele frequencies among eastern UP Population

**Table 2.** Total number of samples classified according to gender

| Sections | No. of O phenotype | No. of A phenotype | No. of B phenotype | No. of AB phenotype | Total |
| --- | --- | --- | --- | --- | --- |
| **Females** | 20 | 17 | 25 | 6 | 68 |
| **Males** | 49 | 24 | 48 | 11 | 132 |
| **Total** | 69 | 41 | 73 | 17 | 200 |

**Table 3.** Rh blood group among eastern UP population

| Phenotypes | Observed |  | Allele frequencies |
| --- | --- | --- | --- |
|  | Number | Percentile |  |
| **Rh(Anti D)+** | 192 | 96% | 0.8 |
| **Rh(Anti D)-** | 8 | 4% | 0.2 |
| **Total** | 200 |  |  |

In India ABO and Rh frequency is studied in other states very well-Karanatak (Prabhakar et al, 2005; Dore Raj and Reddy, 2010), Andhra Pradesh (Naidu et al,2002; Rao et al,2003; Reddy and Reddy,2003, 2005; Reddy and Sudha,2009), Maharastra (Mukherjee et al, 1997; Warghat et al, 2011), Dadar and Nagar Haveli (Meitei and Kshatriya,2010), Rajasthan (Thukral and Bhasin, 1990),Haryana (Kushwaha et al,1990), Punjab (Sidhu, 2003), Uttrakhand (Patni and Yadav, 2003; Guniyal, 2006; Pattanayak,2006), Jammu and Kashmir (Calcutti et al, 2003), Himanchal Pradesh (Mukhopadhyaya and Kshatriys, 2004), Assam (Chakraborty, 2010), Manipur (Devi and Singh, 2008), Orissa (Mohanty and Das, 2010) , Uttar Pradesh (Mandal ,1992; Ara et al 2008; Kumar et al.,2009a,b,c.,2010; Rai and Kumar, 2010,2011, Rai,2011; Rai et al., 2009a,b,c, 2010,), but the UP population is very less explored.

We compared our result with other studies carried out in different parts of the world like-Britain, Guinea, Nigeria, Yamen, Iran, Bahrain, Turkey, Palestine, Saudi Arabia, Pakistan, Nepal and Bangladesh (Mwangin et al,1999; Pramanik and pramanik,2000; Al-Arrayed et al, 2001; Bashwari et al, 2001; Hameed et al,2002; Bahaj ,2003; Boskbady et al, 2005; Dilek et al, 2006; Skaik and El-Zyan,2006; Loua et al, 2007; Firkin et al, 2008; Talukdar and Das,2010). A comparison of frequency of blood group with this study to other is shown in the table 4. Except turkey, Palestine, Nepal and India (present study), frequency of O blood group is highest in Britain (47%), Guinea (48.9%), Nigeria (54.2%), Yamen (51.5%), Iran (34.7%), Bahrain (49.6%), Saudi Arabia (51%), Pakistan (49.6%) and Bangladesh (40.6%) and there is no marked difference in incidence of O blood group in these countries (table 4). Difference is marked in case of A blood group between Turkey and Bangladesh (43.8% vs 26.6%) and is highest phenotype in Turkey (43.8%), Palestine (40%) and Nepal (34%). In Britain B blood group is exceptionally low i.e. 8%, whereas it is highest in our study (36.5%). Marked difference of incidence of AB group is observed between Nigeria and Bangladesh (2.8% vs 9.6%). Rh-negative frequency is lowest in Palestine (2.7%) and highest in Iran (11.3%). Rh-negative frequency in present study is comparable with other countries-Guinea, Nepal and Bangladesh (between 3 to 4%) but is exceptionally high in Britain (17%).

**Table 4.**
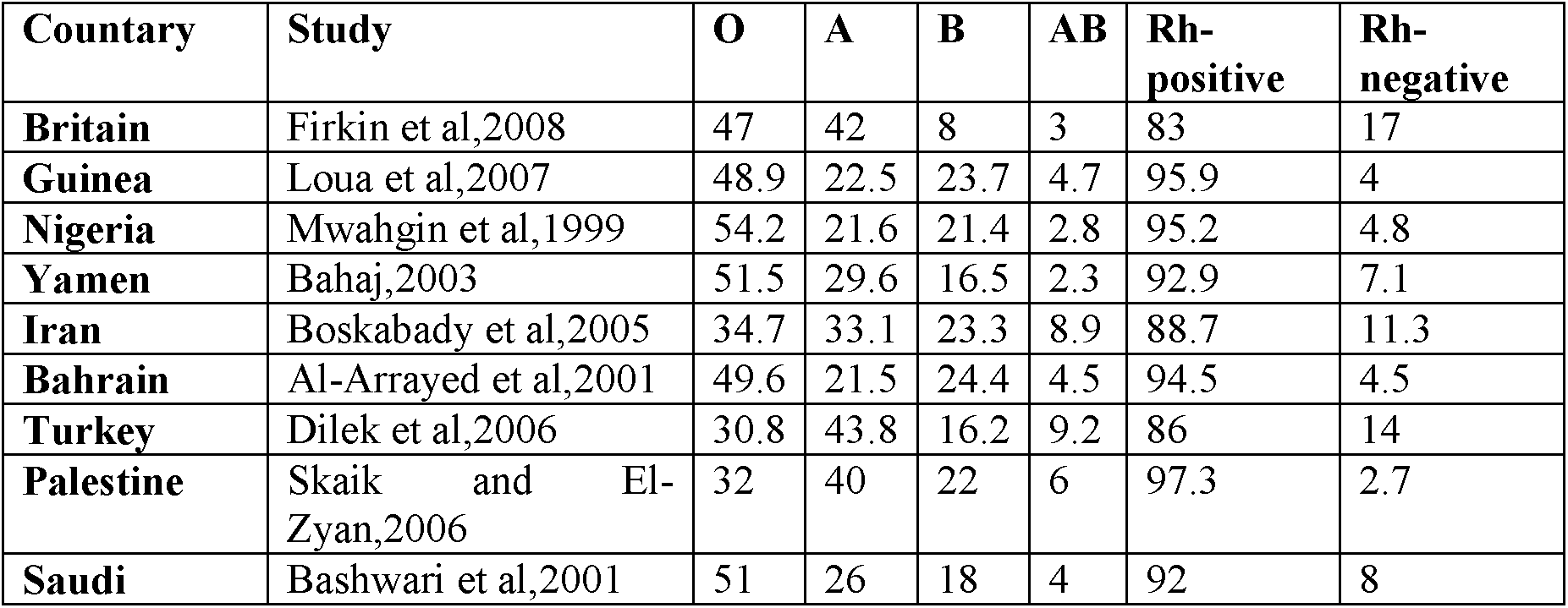
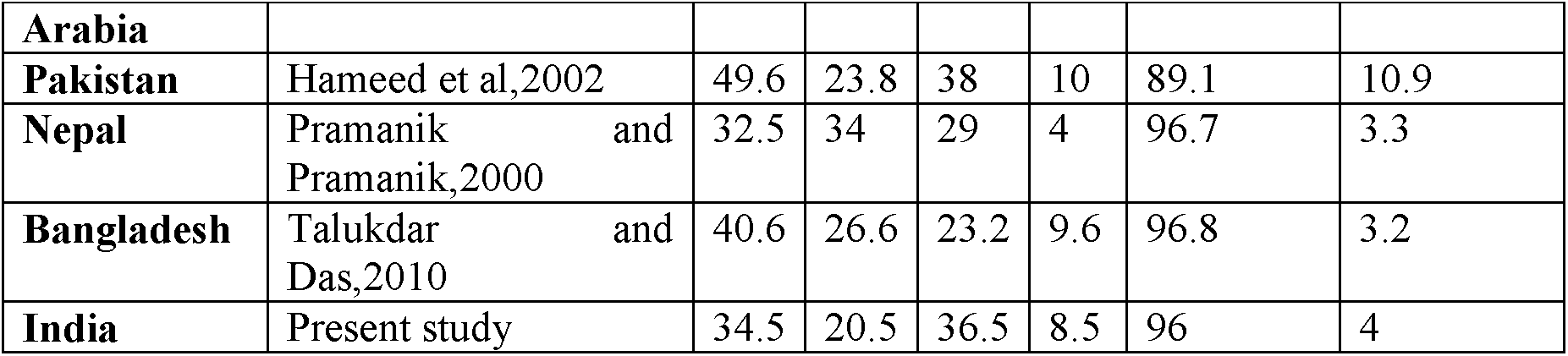
Comparison of percentage frequencies of ABO and Rh blood groups in different studies carried out in different countries

## Acknowledgement

We are grateful to the individuals/subjects who participated in this study and without their cooperation this study could not be completed.

